# Binning microbial genomes using deep learning

**DOI:** 10.1101/490078

**Authors:** Jakob Nybo Nissen, Casper Kaae Sønderby, Jose Juan Almagro Armenteros, Christopher Heje Grønbech, Henrik Bjørn Nielsen, Thomas Nordahl Petersen, Ole Winther, Simon Rasmussen

## Abstract

Identification and reconstruction of microbial species from metagenomics wide genome sequencing data is an important and challenging task. Current existing approaches rely on gene or contig co-abundance information across multiple samples and *k*-mer composition information in the sequences. Here we use recent advances in deep learning to develop an algorithm that uses variational autoencoders to encode co-abundance and compositional information prior to clustering. We show that the deep network is able to integrate these two heterogeneous datasets without any prior knowledge and that our method outperforms existing state-of-the-art by reconstructing 1.8 - 8 times more highly precise and complete genome bins from three different benchmark datasets. Additionally, we apply our method to a gene catalogue of almost 10 million genes and 1,270 samples from the human gut microbiome. Here we are able to cluster 1.3 - 1.8 million extra genes and reconstruct 117 - 246 more highly precise and complete bins of which 70 bins were completely new compared to previous methods. Our method Variational Autoencoders for Metagenomic Binning (VAMB) is freely available at: https://github.com/jakobnissen/vamb

## Introduction

Metagenomics is the study of all genomes in an environment rather than investigating the genomes of single cultured organisms^1^. Current sequencing techniques produce millions of unlabeled reads, each representing a random fragment of one DNA molecule in the sample. To reconstruct the genomes present in the sample, the set of reads are typically reduced to a smaller number of contiguous sequences (*contigs*) by collapsing groups of reads with a significant overlap to the smallest possible superstring in a process called assembly^2,3^. As metagenomes typically contain both highly similar but distinct sequences as well as low-abundance sequences, metagenomic assembly is complex and the majority of observed genomes will appear fragmented in many short contigs. In order to reconstruct genomes from metagenomics wide genome sequencing data, one has to group contigs together by their genome of origin. This task is termed metagenomics binning^4^.

Despite significant advances in both assembly and binning the recent years^5^, binning remains an inaccurate process, where the majority of genomes are fragmented across multiple bins, and most bins are contaminated with sequences from other genomes^6^. Binning tools typically rely on one or a combination of three approaches: *Co-abundance*, which exploits that sequences from the same genome are expected to have a similar abundances across samples, and thus entails grouping contigs with a similar inter-sample abundance profile^7–9^; *composition*, where contigs are grouped by intrinsic properties of the sequences such as GC-content or *k*-mer frequency profiles. These properties have been found to be more similar within phylogenetically closely related contigs than between distantly-related contigs^10–12^. Finally, some methods use *database lookups*, where contigs are grouped based on their similarity to entries in a database of known sequences^13–15^. Unfortunately, because the large majority of existing microbial genomes are not in any database, tools relying on database lookups often face the dilemma of either not assigning a given sequence, or assigning it to distantly related entries, yielding false positives^4^. Examples of binning programs are Canopy^7^ and GroopM^16^ that rely solely on co-abundance information, as well as METABAT^17^, BMC3C^18^, CONCOCT^8^ and MaxBin^19^ which rely on both co-abundance and compositional information.

Deep learning is a field in massive expansion with great successes in image and speech recognition, natural language understanding, computational biology and artificial intelligence for games^20,21^. An autoencoder^22^ consists of two neural networks, the *encoder*, which maps a high-dimensional input to a low-dimensional encoding called the *latent representation*, and a *decoder*, which maps the encoding to an output of same dimensions as the original input. The autoencoder is trained to minimize the difference between the input and output, and if successful, necessarily learns to represent the information of the input in a lossy but lower-dimensional encoding. A variational autoencoder (VAE) is a generative model where the latent representation is stochastic, for example Gaussian, and thus encoded by a mean and a standard deviation for each component^23,24^. The VAE is relevant here because it allows an easy implementation of a constraint on the information in the model, which prevents the VAE from learning to simply copy the input. In the VAE framework, this is done by penalizing the Kullback-Leibler (KL) divergence between the latent posterior distribution defined by the encoder and the (Gaussian) latent prior distribution. VAEs are able to identify co-variation patterns in data beyond linear relations. VAEs have been explored in many aspects such as unsupervised learning^25^, semi-supervised learning^26^ and supervised learning^27^ as well as many different architectures such as the ladder VAE^28^, beta-VAE^29^, Info-VAE^30^ and others. Each contig is represented as a point in a high dimensional space where the dimensions express its *k*-mer composition and per-sample abundance. Hereafter, binning of the contigs is simply the clustering of points in high-dimensional space. In this view, we use VAEs to denoise the input and increase the separation between the clusters. Furthermore, we hypothesized that the network would efficiently learn to integrate the different data types of abundance and sequence composition and through these two mechanisms improve binning.

In this study, we trained a VAE to encode abundance and composition data from metagenomics samples. We clustered the resulting latent representation using an iterative medoid clustering algorithm and using three different benchmark datasets we show that our approach produced 1.8 - 8 times more highly precise and complete bins sets compared to existing state-of-the-art. In addition, we demonstrate the ability of our approach to encode and cluster large gene catalogues, where we cluster 39-70% more genes and reconstruct 51-58% more large bins compared to Canopy and MSPminer on the human gut microbiome Integrated Gene Catalogue (IGC). Furthermore, our approach produced 21-57% more highly precise and complete bins from the IGC dataset compared to the other two binners and of which 70 of the bins were exclusive to VAMB. Our study also demonstrates the power of the variational autoencoders to easily integrate heteromorphic data. This is a common issue in biology today where cells, samples and individuals are often assayed with multiple different technologies such as genomics, transcriptomics and proteomics. Our results show that this can be done with co-abundance and sequence data without developing statistical models for each data type. The method, Variational Autoencoders for Metagenomic Binning (VAMB), is licensed under MIT license and freely available at https://github.com/jakobnissen/vamb.

## Results

### Using a VAE for binning metagenomics data

The input to the VAMB pipeline is a catalogue of metagenomic sequences that needs to be binned, representing either the assembled DNA fragments or genes of the metagenome, and their abundances. Usually, such a contig catalogue can be obtained by per sample *de novo* assembly. Alternatively, gene catalogues, such as the Integrated Gene Catalogue (IGC)^31^ of the human gut microbiome, can be produced by predicting coding sequences in the metagenomic assembly, and clustering the genes by sequence identity. In both cases, the associated sequence abundance can be obtained by mapping reads to the sequence catalogue. The VAMB pipeline consists of three major steps (Figure 1).

**Figure 1.**
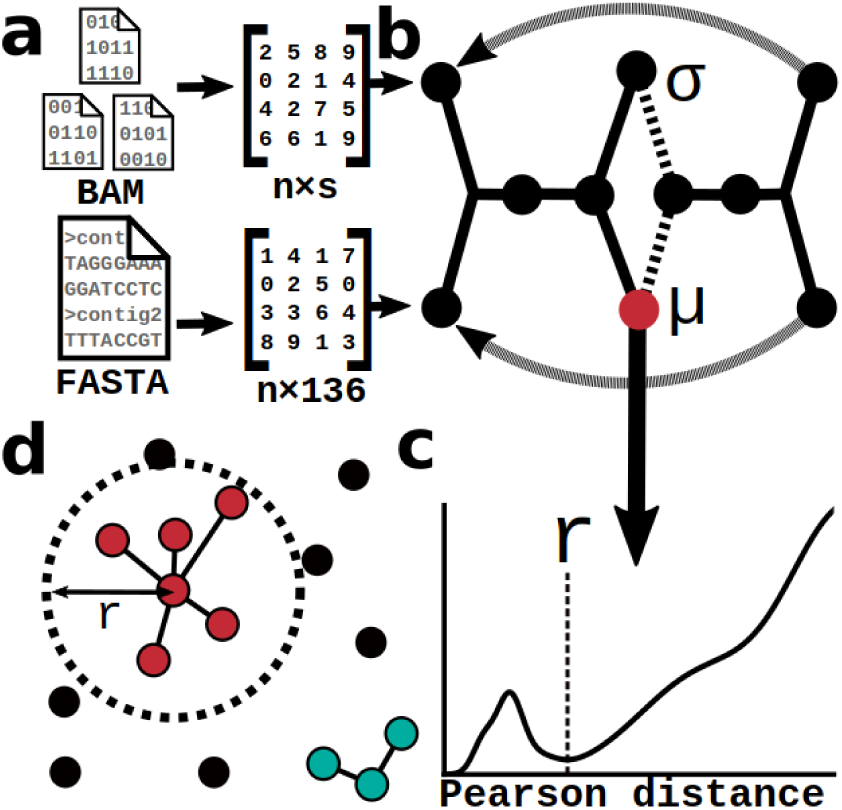
Overview of the VAMB workflow. **a**) Tetranucleotide frequencies (TNF) from gene or contig sequences were calculated from the sequences, and the abundances were determined from BAM files with read mappings to the gene or contig dataset. **b**) The variational autoencoder was trained for 500 epochs with the TNF and abundance tables as input. Hereafter the data was passed through the encoder and the mean of the latent neurons were extracted and used as a latent representation of the input data. **c**) The density of pearson correlation coefficient distances between latent representation of genes or contigs separated in small groups of similar sequences and a large group of distant sequences. The clustering threshold (r) was determined as the minimum between the two groups. **d**) Using iterative medoid clustering, all genes or contigs within the pearson distance threshold (r) of the medoid contigs were clustered as a bin.

First, for each DNA sequence in the gene or contig catalogue, the per-sequence Tetra-Nucleotide Frequencies (TNF) for the 136 possible canonical 4-mers is calculated and the abundance of each sequence is estimated based on read mappings. In the second step, these tables are concatenated and used to train a VAE *tabula rasa*. The VAE passes the concatenated data through two hidden layers and encodes it as a multivariate Gaussian latent distribution (Figure S1). To reconstruct data in the VAE we sample the latent distribution and pass this representation through the decoder with two hidden layers and an output layer with the same dimensions as the input layer. The reconstruction loss consists of a cross entropy (CE) loss of the abundance reconstruction as these can be interpreted as a probability distribution and the sum squared errors (SSE) loss of the TNF. The original VAE loss defines a lower bound on the marginal likelihood of the input data known as the evidence lower bound (ELBO). Since our objective is clustering and not the marginal likelihood we introduced hyperparameters weighting the three loss terms CE, SSE and KL divergence. Usually downweighting the KL will give better clustering because the KL term penalizes deviation from a Gaussian distribution. After training, the DNA sequences and co-abundance information of the gene or contig catalogue are encoded to the mean of their latent distributions. Last, the latent representation is randomly sampled to estimate a threshold for clustering based on Pearson distance between contigs, before iterative medoid clustering is applied to the encoding to construct the final bins.

### Hyperparameter search for the VAE

To identify the best hyperparameters to use in the VAE we trained and benchmarked using two different training datasets, and used a third held out dataset as validation. As one training dataset, we used the high complexity dataset from the first Critical Assessment of Metagenome Interpretation (CAMI)^32^ consisting of five simulated samples in a time series with a total of 596 genomes and 478 circular elements. Because the simulated contigs had a length distribution that was unrealistic compared to that of real metagenomes (Figure S2), we reassembled the data using metaSPAdes^3^ to 275,285 contigs and processed this dataset (CAMI High). The second training dataset was the MetaHIT “error-free” dataset originally created by Kang and co-workers^17^ that used 290 sequenced genomes from the human gut microbiota shredded into 182,388 contigs as a contig catalogue for 264 human gut sequencing runs^33^. Our validation dataset, CAMI Airways, was the simulated Illumina sequences of the ten human airways samples from the 2nd CAMI challenge human toy dataset, comprising 187,685 contigs from 639 OTUs.

We performed two rounds of hyperparameter search on the two training datasets. To evaluate the hyperparameters, we measured performance in terms of the number of true bins reconstructed with per-nucleotide precision above 0.9 and at a range of recall thresholds (0.3 - 0.95). In the first round, we trained and evaluated 50 networks for each dataset, where each hyperparameter was chosen randomly from an interval (Figure S3). Among the tested hyperparameters, we found that dropout, warmup, and the loss scaling parameters α (sseratio) and β (errorsum) impacted performance. In contrast, the number of training epochs and the number of latent neurons seemed to have little effect. For all hyperparameters, the variation in performance due to confounding hyperparameters was large and we therefore selected a tentative value for each. The we performed a second round of evaluation, where each hyperparameter varied, while the others remained fixed (Figure S4 and S5). The best-performing value was then chosen as the final default value. For a complete overview of initial sampling intervals, tentative values and final values, see Table S1.

### The VAE encodes and integrates metagenomics data

We investigated how autoencoding the input data affected the quality of the genome bins produced. To that end, we compared the bins produced by clustering the raw input data to the bins produced by clustering the latent encoding. When assessing bins, we define “precise bins” as bins at or above a per-base precision threshold of 0.9, and “complete bins” as bins with a per-base recall of 0.9 or more. For all three datasets, we found that the number of precise and complete bins increased when clustering the latent representation compared to the raw data, from 6 to 46 bins for our holdout dataset. We found that this increase was independent of using only TNF, only abundances or using both abundance and TNF as input. The only exception was when using only TNF on the CAMI High dataset, where encoding modestly decreased performance (Figure 2C, Figure S6, Table S2). We observed that clustering the abundance data from the MetaHIT dataset provided more precise and complete genome bins compared to using TNF only (Figure S6).

**Figure 2.**
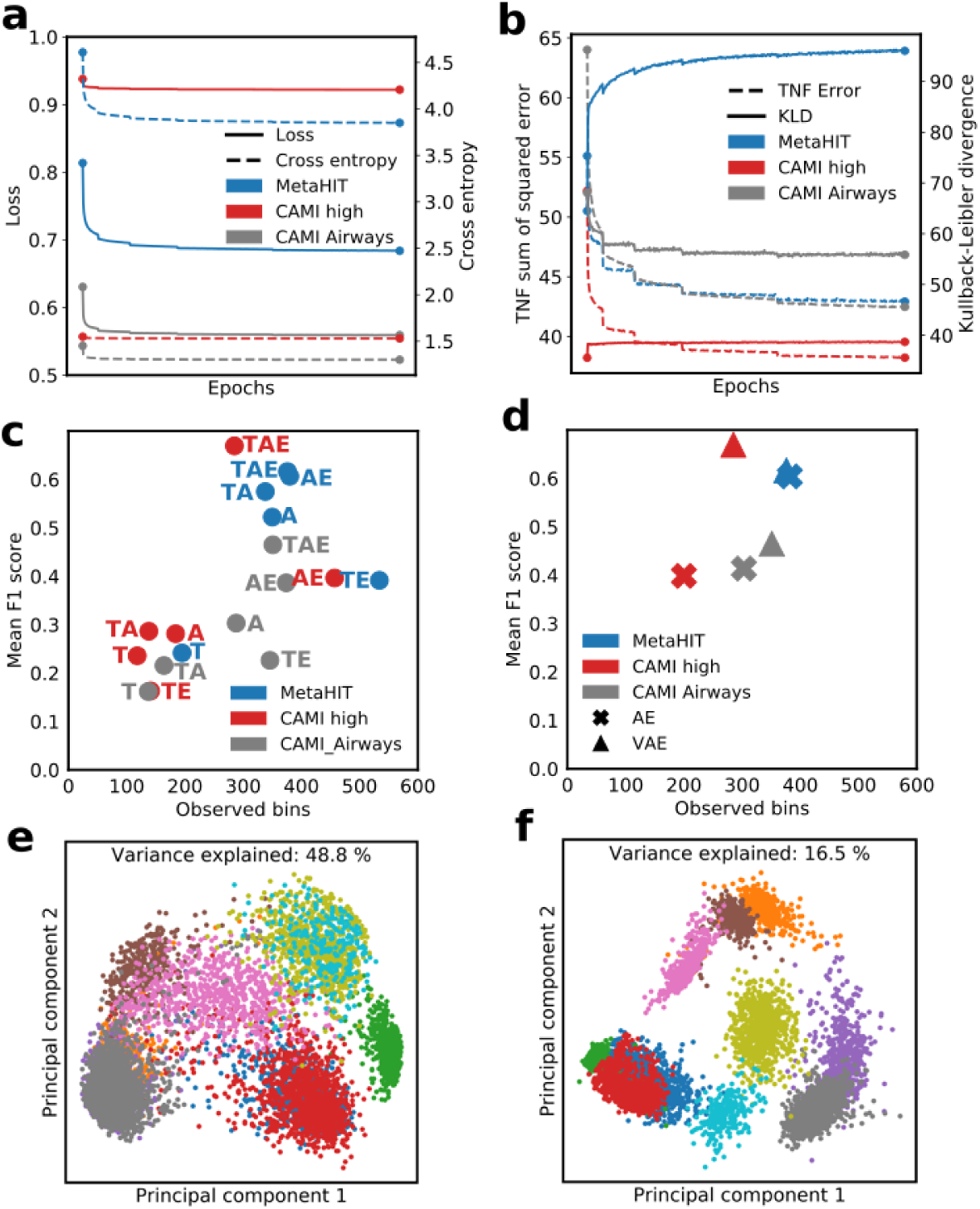
Properties of autoencoding the input data for VAMB. **a**) Loss (solid lines) and abundance cross entropy (dashed lines) during training of the VAE on the three benchmark datasets. Blue: MetaHIT, red: CAMI High, gray: CAMI Airways. **b**) TNF reconstruction as the sum of squared error (dashed lines) and Kullback-Leibler divergence (solid lines) during training of the VAE on the three datasets. Colors as in a. **c**) The effect on performance of clustering different inputs; T = TNF data, A = Abundance data, E = Inputs encoded using the VAE. The F1-score increases when using the latent representation of the VAE. Colors as in a. **d**) Performance when using a VAE compared to an ordinary autoencoder (AE). The VAE (triangle) improves performance over the standard autoencoder (cross), except on the MetaHIT dataset. Colors as in a. **e**) and **f**) Contigs from the same 10 randomly chosen bins of CAMI Airways projected onto their two principal component axes, each bin represented by a color. In **e)**, projection of abundance and TNF data for the ten bins i.e. the input data for VAMB, in **f)**, projection of the latent encoding of the input data. In the latent encoding, clusters are better separated, and, despite the latent encoding having lower dimensionality, the two principal components explain less of the variance.

Interestingly this was the opposite for the CAMI High dataset where clustering was most efficient using only the TNF data compared to using only abundances. This was in line with our expectations due to the different number of samples (5 in CAMI High vs. 124 in MetaHIT). The performance improvement of encoding showed that the VAE was able to make use of the information in TNF data, abundance data or both without any explicit statistical model of how this should be done. To investigate the vulnerability to overfitting, we ran VAMB on the training datasets for up to 2500 epochs and assessed performance, compared to the default training of 500 epochs (Figure S7). Beyond 500 epochs, we saw little change in the performance, implying that the network was resistant to overfitting for at least the first 2500 epochs.

We additionally tested a version of VAMB with the VAE replaced by an otherwise identical ordinary autoencoder and saw significantly degraded performance (Figure 2D, Figure S8) for two of the three datasets, where the number of high-quality bins recovered from our holdout dataset decreased from 46 to 22. Furhtermore, when visualizing the two principal components of the input and latent encoding of the validation dataset, we found that the latent encoding seemed denoised and the clusters separated more clearly (Figures 2E, 2F), in line with the findings of a previous study using a VAE to enhance clustering^34^. We further note that the variance explained by the two principal components decreased from 48.8% to 16.5% after encoding, implying that encoding spreads the data points more evenly across available dimensions.

### VAMB outperforms other binners on bacterial sized genomes

To evaluate the performance of VAMB we compared our approach with three other binners: Canopy, METABAT2 and MaxBin. Following earlier work^17^, we considered the number of precise bins reconstructed at recalls in tiers from 0.5 - 0.99. We found that VAMB reconstructed more precise and complete bins compared to all other binners (Figure 3A, Table S3). The only recall threshold where any other binner produced more precise bins was the lowest recall thresholds 0.5 and 0.6 where METABAT2 produced more bins on the validation dataset, CAMI Airways (Figure 3B). Compared to MaxBin and Canopy, VAMB was far superior and able to reconstruct more bins across all precision and recall thresholds (Figure 3B, Table S3). We found that VAMB reconstructed 24, 42 and 46 precise and complete bins for the MetaHIT, CAMI High and CAMI Airways datasets, respectively (Table 1). In comparison, the next best binner, METABAT2, reconstructed 3, 16 and 26 for the same datasets, therefore, VAMB reconstructed between 1.8 - 8 more precise and complete bins. Compared to Canopy and MaxBin, VAMB reconstructed between 8 - 46 and 4.6 - 42 more precise and complete bins, respectively (Figure 3B, Table 1). Importantly, as we optimized the hyperparameters of the VAE network based on the MetaHIT and CAMI High datasets, we used the CAMI Airways dataset as an independent assessment of our default parameters. Here, VAMB outperformed all the other binners at high recall thresholds and reconstructed 11 genomes with a precision and recall greater than 0.99.

**Table 1.** Number of bins created by the different binners for the three different datasets at a precision greater than 0.9 and recall greater than 0.9. For other combinations of precision and recall see Table S2.

| p>0.9 / r>0.9 | Canopy | MaxBin | METABAT2 | VAMB |
| --- | --- | --- | --- | --- |
| MetaHIT | 3 | 1 | 3 | **24** |
| CAMI High | 0 | 1 | 16 | **42** |
| CAMI Airways | 1 | 10 | 26 | **46** |

**Figure 3.**
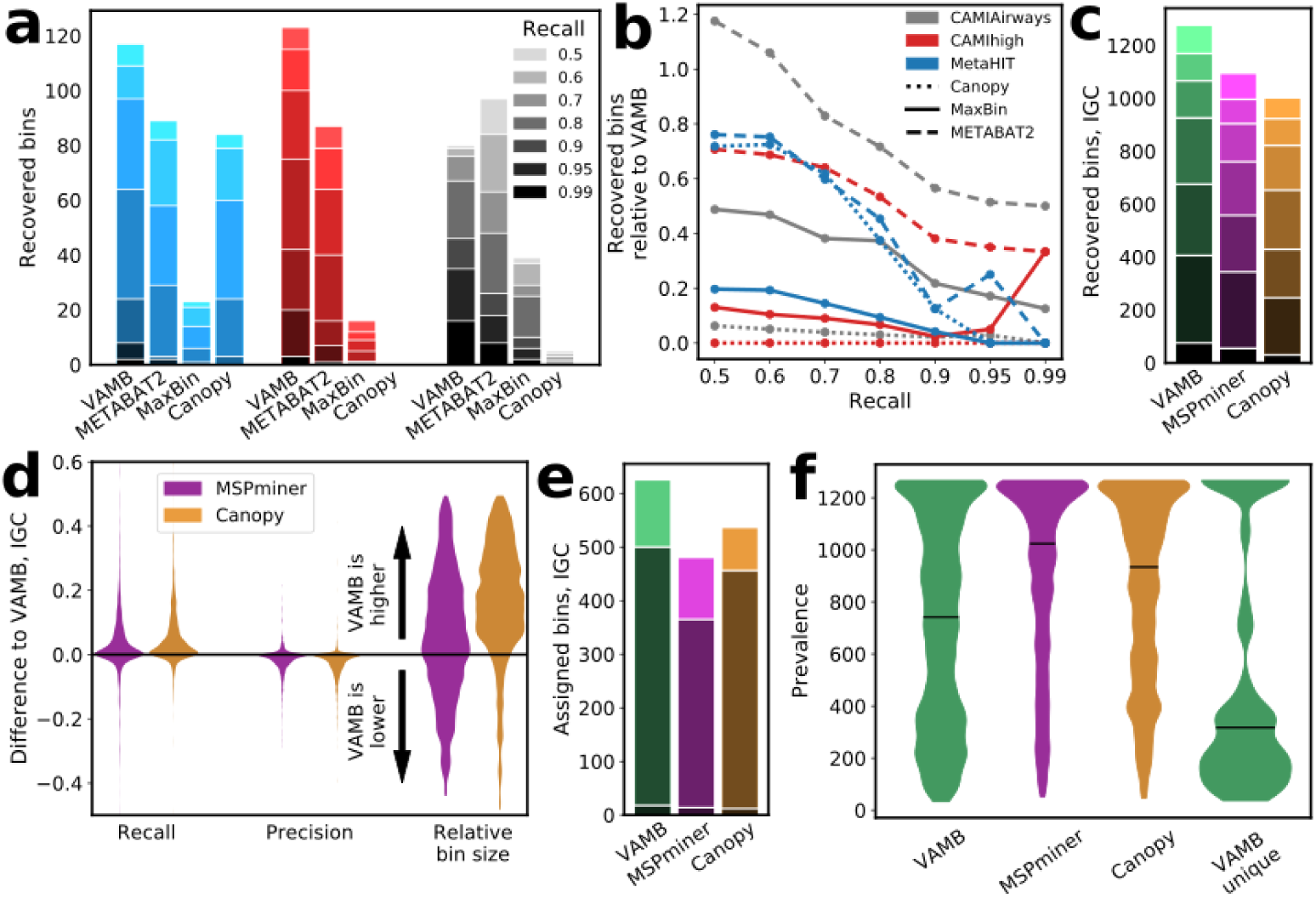
Performance of VAMB compared to other binning tools. The performance was determined using the thresholds of precision ≥ 0.9 and a range of recall thresholds (0.5, 0.6, 0.7, 0.8, 0.9, 0.95, 0.99). **a)** Number of recovered bins with total size ≥ 200 kbp for VAMB, METABAT2, MaxBin2.0 and Canopy for the range of recall values on the MetaHIT (blue), CAMI High (red) and CAMI Airways (gray) datasets. VAMB recovers more large, high quality bins compared to the other binners. Shading of the bars represent recall threshold from light (0.5) to dark (0.99). **b)** Same as a, but expressed as a ratio of the number of bins recovered by VAMB. As recall increases, the relative gain of using VAMB increases. Blue: MetaHIT, red: CAMI High, gray: CAMI Airways, dotted line: Canopy, filled line: MaxBin, dashed line: METABAT2. **c)** Number of recovered large (≥ 500 genes) bins from the IGC dataset at the range of recall thresholds (0.5 - 0.99, light to dark shade). VAMB identified more precise and complete bins compared to the other binners. Green: VAMB, purple: MSPminer, brown: Canopy. **d**) Difference in recall, precision and relative bin size for large (≥ 500 genes) bins that were common between VAMB and one of the binners (MSPminer, Canopy). We defined common bins of two binners as those with a reciprocal overlap of ≥ 50%. The IGC bins created by VAMB tend to have a higher recall and larger number of genes (size), but lower precision. **e**) The number of large bins with recall ≥ 0.5 which can be assigned to genus (light color), species (darker color) or subspecies (darkest, near zero) based on alignment to databases. Green: VAMB, purple: MSPminer, brown: Canopy. **f**) Distributions of the prevalence, defined as number of samples with a nonzero abundance for precise and complete (precision and recall ≥ 0.9) IGC bins. The horizontal line represents the median. Shown for each binner, plus the bins uniquely found by VAMB. VAMB is able to reconstruct bins present in fewer samples. All violin plots were smoothed with a gaussian kernel with bandwidth of 0.1, and the width of the violins have been scaled by their widest point.

### Application on large datasets using gene catalogues

When analyzing large-scale datasets with hundreds to thousands of samples, genes are often used as the sequence unit in non-redundant gene catalogues. This is because contigs can be large and have no natural beginning or end making homology reduction much more complicated^31,33^. However, clustering catalogues with millions of genes and thousands of samples are computationally intensive^31,35^. When we instead encode the gene catalogue to a latent representation of smaller dimensionality, clustering is faster.

We therefore used VAMB to train on and encode the Integrated Gene Catalogue (IGC) of 9.8 million genes and 1,270 samples into a latent representation that could be clustered. We used a single Graphical Processing Unit (GPU) to train the VAE on the abundance and TNF data and found the TNF loss to increase after a few epochs. This indicated that the latent space of only 40 neurons was not enough to contain both abundance and sequence information and the network prioritized reconstructing the abundance rather than the TNF (Figure S9). This makes sense as there was a large number of samples (1,270) combined with short sequences (genes) which increased the noise in the TNF data. We, therefore, increased the number of latent neurons to 80 and removed dropout which made the model train as expected (Figure S9). Hereafter, we clustered the latent representation sampling out 30,000 bins. In total, 5.5 million genes were clustered after which the number of new large bins was small (Figure S10). When filtering bins for a size of 500 or more genes a total of 4.60 million genes were clustered in 2,378 bins (Table S4). This represents 1.29 million (39%) more genes in 876 more bins (58%) compared to MSPminer^36^ that clustered 3.31 million genes into 1,502 large bins. Compared to Canopy this was 1.89 million more genes (70%) and 800 (51%) more large bins. Because the ground truth is unknown for the IGC dataset, we estimated performance using known marker gene sets^37^. Here we were able to identify 675 precise and complete bins, an improvement of 117 and 246 compared to MSPminer and Canopy that reconstructed 558 and 429 bins at the same thresholds, respectively. At lower recall thresholds (0.5, 0.6, 0.7 and 0.8), the improvement of VAMB compared to MSPminer and Canopy was consistent, as VAMB reconstructed 162 - 182 and 244 - 275 more precise bins, respectively, depending on the threshold chosen (Figure 3C, Table S4).

We investigated the set of bins that were common between the methods (defined by 50% reciprocal overlap of bin content) and found that the common bins created by VAMB were generally larger than their counterparts from other binners (Figure 3D and Table S5). Compared to the bins produced by MSPminer and Canopy, the bins created by VAMB had, on average, 53 and 405 more genes, respectively. This resulted in a higher recall with an average increase of 0.034 and 0.057, but also slightly lower average precision of -0.017 and -0.027 (Figure 3D). One cause of the lower precision of VAMB compared to the other binners could be that VAMB combined genes from different bacterial strains into one bin. To test this, we allowed strain heterogeneity within a bin as determined by CheckM^37^ and found that VAMB reconstructed relatively more precise and complete bins (809 compared to 616 and 431 for MSPminer and Canopy, respectively) and that the average decrease in precision was lowered to -0.0007 and -0.0025.

In addition to the marker gene analysis, we determined the taxonomical consistency of bins based on alignment to bacterial genomes. In general we found that more bins from VAMB could be annotated (Figure S11) and that this was also true for the lower (i.e. more specific) taxonomic ranks. Here VAMB, MSPminer and Canopy produced 624, 479 and 535 bins that could be annotated to genus or lower rank (Figure 3E), respectively. Additionally, for the overlapping bins we found a high consistency between the binners, where 77% of the common bins were annotated to the same taxonomy.

Finally, we investigated the precise and complete bins that were not common bins, i.e. unique to each binner. Here VAMB, MSPminer and Canopy produced 70, 4 and 4 unique, precise and complete bins, respectively. When investigating the prevalence of the precise and complete bins we found that VAMB in general reconstructed more relatively rare bins and that the unique bins were divided in two distributions (Figure 3F). A minor set of the bins (n=16) were present across all samples, whereas another subset (n=48) was primarily present in a small proportion samples of up to approximately 500 individuals. Furthermore, among the 70 unique bins from VAMB, we identified 9 that could be taxonomically annotated to genus or species level. Among those, and of particular taxonomic purity were bins assigned as *Elusimicrobium sp. An273* and *Desulfovibrio sp. An276*, two anaerobic species that have recently been identified in the chicken gut microbiome^38^. The remaining 61 bins uniquely found by VAMB that could not be annotated at genus or species level, despite being precise and complete, potentially represent novel microorganisms not previously described from the human gut microbiome and the IGC dataset.

### VAMB can be run on standard computational infrastructure

When developing VAMB we were interested in creating a new binning tool that based on deep learning would be available to most users without access to a high-performance computational system. We therefore developed and tested the binners on a single Linux machine with an Intel Xeon E5-4610 CPU using 4 virtual cores. When running the binners on the contig datasets we found the runtime of VAMB to be slightly longer (0.96 - 1.6 times) compared to METABAT2, whereas memory usage was lower at 0.24 to 0.44 times at much (Table 2). All contig datasets were trained and clustered using VAMB in less than 3 hours and using less than 1 Gb RAM. For the IGC dataset, we used a single GPU to accelerate the training and was able to train the VAE in approximately 48 hours. Hereafter, we clustered the latent representation using 16 CPU cores and 6 Gb RAM to create 30,000 bins in approximately 5 days. We note that it is possible to train the network using CPUs. In comparison we clustered the IGC dataset using Canopy using 32 CPU cores for 7 days using 190 Gb RAM, however Canopy might have plateaued earlier.

**Table 2.** Computational resources needed to run the different pipelines for the four different binners on the three different contig datasets and the Integrated Gene Catalogue (IGC). *: the VAE was trained using a Tesla Quadro M6000 GPU and clustered on a CPU. Canopy was allowed to perform clustering for 7 days on the IGC dataset, before clusters were reported. For IGC the RAM usage for VAMB was 7 Gb for clustering and 59 Gb for encoding, and 7200 mins for clustering and 2880 minutes for training the VAE.

|  | MetaHIT |  | CAMI High |  | CAMI Airways |  | IGC* |  |
| --- | --- | --- | --- | --- | --- | --- | --- | --- |
|  | Time (min) | RAM (GB) | Time (min) | RAM (GB) | Time (min) | RAM (GB) | Time (min) | RAM (GB) |
| Canopy | 1 | 0.19 | 1 | 0.18 | 2 | 0.37 | 11,500 | 189 |
| MaxBin | 847 | 2.64 | 348 | 3.12 | 1459 | 3.50 | NA | NA |
| METABAT 2 | 139 | 2.12 | 142 | 2.10 | 75 | 2.29 | NA | NA |
| VAMB | 134 | 0.94 | 177 | 0.75 | 121 | 0.55 | 7200 (2880) | 7 (59) |

### Discussion

Binning of assembly data is an important step in metagenomics studies as it reduces the complexity from often millions of contigs or genes to hundreds or a few thousand microbial species^7^. For studies of the human microbiome, this enables association analysis between microbial species, genetic entities such as plasmids and phages with host, clinical and cohort information^39^. For metagenomics studies of unknown microbial communities, binning allows the reconstruction of genomes from microorganisms in the environment. This enables an initial description of the community and can be a starting point for further analysis of the organisms. Here, we provide the first attempt at combining metagenomics binning with unsupervised deep learning and show improvement compared to state-of-the-art methods. We found that our framework works well across datasets of different types and sizes. For the contig datasets VAMB reconstructed several times (1.8 - 8) more precise and complete bins compared to the other methods and for the gene-catalogue dataset an extra 117 - 246 precise and complete bins. Additionally, VAMB was able to reconstruct bins that were less prevalent compared to the other binners.

Benchmarking of metagenomics binning is not a straightforward task, even when the true bins are known, and several efforts have been initiated to address this^32^. For example, there is no consensus on what constitutes a good set of bins, as some situations call for reconstructing few but high-quality genomes, whereas other situations may weigh reconstructing as many of the present genomes as possible within some lax constraints on precision. Additionally, it is unclear how to deal with the microdiversity of very similar strains typically found in metagenomes. In our case, we have designed VAMB to be highly performant according to the criteria set by the Critical Assessment of Metagenome Interpretation (CAMI), namely the number of reconstructed high-quality (high recall and precision) genomes^32^. Because our assessment of the IGC dataset is based on marker gene sets, we additionally assessed the performance based on taxonomic assignments inferred from alignment of genes to sequence databases. Here we verified the increased performance of VAMB showing a more bins to have specific taxonomic assignment across all levels.

We believe that the increased performance of VAMB compared to the other binners is mainly achieved by the VAE learning the manifolds of the data that corresponds to the microbial entities in assembly space. This will force the co-abundance and TNF profiles of noisy genes or contigs to be represented similarly to the manifold of the main cluster and thereby reduce noise in the data. Additionally, we show that two datasets of observations, here co-abundance and TNF, can be integrated by the deep model and that the resulting latent representation is more informative than either of the inputs. Interestingly, this is in principle not limited to only two input data types and it is possible to add additional data as input to the VAE. For example, Yu and co-workers^18^ have recently demonstrated that adding codon usage of contigs provides information not contained in the TNF, and improves binning^18^. Modifying VAMB to use more inputs such as codon usage or other information would be straightforward.

When developing our model we investigated how changing the architecture of the VAE as well as various modifications to the loss function could affect the clustering performance. We found this to be of particular importance as an improvement in the loss of the network did not necessarily lead to improved clustering of the latent representation. Other implementations of the VAE such as e.g. the Info-VAE^30^ was much slower to train and did in our case not lead to a rich latent representation. With regard to using VAEs for clustering of unstructured data into microbial species, this problem is somewhat analogous to clustering of single-cell RNA data by cell populations. Here more complex models such as e.g. Gaussian Mixture VAEs (GM-VAE) has been applied where the cell clustering is an output of the VAE model itself^25,40^. However, when the number of clusters for the GM-VAE reaches hundreds or thousands, as it common within a microbial community, this approach becomes computationally infeasible in most settings. In addition to adding other datasets as input to VAMB, we envisage that one could develop convolutional layers to learn the sequence information from the sequences themselves. Furthermore one could develop and benchmark approaches based on generative adversarial networks^41^ as described in Ghahramani et al. for scRNA^42^.

Finally, we believe that the importance of our findings is not limited to the fields of microbiome and metagenomics as data integration is a central process in many fields of life science research. Current and future discoveries within fields such as genomics, microbiome, cancers genomics and precisions medicine will be greatly enhanced by data integration across several omics datasets as well as clinical information. To achieve this, deep learning methods, such as the VAE or other models provide promising approaches.

## Methods

### Computation of abundance and tetranucleotide frequencies

For each contig, the frequencies of each 4-mer not containing ambiguous bases were calculated to obtain TNFs. Because the coding strand was unknown, the TNFs were grouped with their reverse complement, resulting in 136 canonical TNFs. Thus, for *n* contigs, the output was an *n* × 136 table. To determine abundance, we counted the number of individual reads mapped to each contig with a mapping score above 50. Alternative mapping hits marked with “XA” in the BAM files were counted on equal footing with primary hits. Each read in a read pair was counted independently, with a read counted twice if its mate was unmapped. The read counts were normalized by contig length and total number of mapped reads, such that abundance was given in reads per kilobase contig per million mapped reads (RPKM). With *s* samples and *n* contigs, the abundance output was an *n* × *s* table. Abundance values were normalized across samples to sum to one in order to mimic a probability distribution that was reconstructed from the final VAE by applying softmax to the abundance output neurons. Finally, TNFs was normalized by z-scaling each tetranucleotide across the contigs in order to increase the relative inter-contig variance.

### Architecture of the variational autoencoder

Each contig was input to the VAE as an abundance vector A_in_ of length *s* and a TNF vector T_in_ of length 136. These were concatenated to a vector of length *s* + 136, before being passed through the hidden encoding layers consisting of two fully-connected layers, each using batch normalization^43^ and dropout^44^ (p = 0.2). The output of the last layer was passed to two different fully-connected layers of length N_L_, termed the µ and the σ layer. The latent layer, *l* is a vector of length N_L_ obtained by sampling the Gaussian distribution: *l*_i_ ∼ N(µ_i_, _i_) for each neuron *i* = 1…N_L_. The sampled latent representation was then passed through the hidden decoding layers, identical in size to the hidden encoding layers, except arranged in reversed order. Finally, the last hidden decoding layer was connected to a *s* + 136 fully connected layer, which was split into two output vectors A_out_ and T_out_ of length *s* and 136, respectively. After training, the input sequences were encoded by passing it through the VAE and extracting the values of the µ layer.

The VAE models were trained using the Adam optimizer^45^ and using one Monte Carlo sample of the Gaussian latent representation. For the contig catalogues (MetaHIT, CAMI High and CAMI Airways) we used a mini-batch size of 64 that doubled after 25, 75, 150 and 300 epochs to a final of 1024, a learning rate of 10^-3^ and trained for 500 epochs. For the IGC dataset, the network was trained by setting a starting mini-batch size of 128 which increased to 2048 at the same epochs as specified for the contig datasets, a learning rate of 10^-5^, and a dropout probability of 0. The VAE was implemented using PyTorch^46^ v 0.4.1 and when using a GPU running CUDA (v.8.0.61).

### Loss function

When training the VAE with *s* samples and *N*_*L*_ hidden neurons, the failure to reconstruct the input was penalized by the reconstruction error, consisting of an abundance error (E_ab_) and a TNF error (E_TNF_), defined as

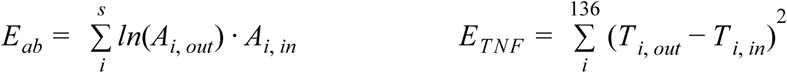

To regularize the model, the *µ* and *σ* layer was constrained by a prior N(0,1) by penalizing the deviance from this distribution with the Kullback-Leibler divergence

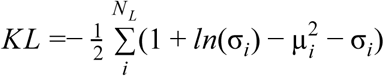

Finally, the combined model loss was then:

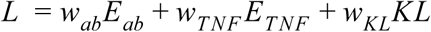

where the weighting terms are defined as *w*_*ab*_ = (1 – α) *ln*(*s*) ^−1^, *w*_*TNF*_ = α/136 and *w*_*KL*_ = (*N*_*L*_β) ^−1^. The parameters α and β were set to 0.05 and 200, respectively.

### Determination of clustering threshold

A number of seed contigs were randomly sampled from the latent encoding. The Pearson distances of the seed contig *s* to every other contig *c*, 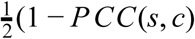, where PCC(*s, c*) is the Pearson correlation coefficient, were calculated and binned to 400 equidistant bins in the interval [0, 1]. The density of binned distances was smoothed using a Gaussian kernel *N*(0, 0.015), and numerically differentiated. The threshold between the close and far contigs was defined as the first minimum after the first local maximum, where fewer than 2,500 contigs were closer than the threshold. For the training the IGC dataset, we manually set this to 10,000 genes to control for the higher amount of genes compared to contigs in a typical microorganism. If no threshold was found, the threshold was set to the first point where the density was 0.8 times the maximum and where fewer than 2,500 contigs or 10,000 genes were closer than the threshold. If still no threshold was found, the threshold was undefined. This process was repeated 2,500 times, and the median threshold chosen for the clustering of the latent representation.

### Clustering

Clustering of the latent space was done using an iterative medoid clustering approach (Supplementary Information Algorithm S1). In our clustering algorithm, a *seed contig* is arbitrarily chosen, and the *inner contigs* defined as all other points within a certain (Pearson) distance from the seed. Each of the inner contigs was then randomly sampled and accepted as seeds if the mean distance of their inner points were less than the previous seed’s mean distance. After 15 consecutive futile samples or if all inner contigs have been sampled, the inner contigs were returned as a cluster and removed from the dataset. A new seed was chosen and the process continued until all points have been clustered or the specified maximum number of seeds were reached.

### Assigning ground truth to the reassembled CAMI High dataset

To asses which reassembled contigs truly belonged to which bin, we aligned the reassembled contigs to the original using blastn^47^ (v. 2.2.31). We removed any hits shorter than 500 bp or with lower nucleotide identity than 95%. If a query (reassembled) contig was aligned to multiple reference (original) contigs, we accepted the reference with the longest alignment, if the alignment was more than twice as long of the next longest. If that was not the case for any reference, we accepted the reference with highest nucleotide identity, if the reference was longer than 10 kbp, had an alignment length of at least 90% of the longest-aligning reference, and had at least 0.05% higher nucleotide identity than the second-highest identity reference. If no reference fit those criteria, they were ignored in the benchmarking.

### Benchmarking

When benchmarking a set of bins against a true set of bins, we matched each true bin with each of the observed bins and for each pair, calculated the *precision* as the total size of contigs in the observed bin that belonged to the true bin divided by the total size of the observed bin. The *recall* was defined likewise, but with the size of the true bin in the denominator. Given a threshold precision and recall pair, we counted the number of true bins that matched or exceeded the particular threshold. The F1-score was calculated as.

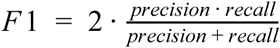

When comparing the performance of VAMB with other binners, we used Canopy from the original published version in 2014 and ran it with default parameters. METABAT (v.2.10.2) was run with default parameters, except setting --minClsSize=1, in order for it to not discard small but accurate bins. MaxBin (v.2.2.4) was run with default parameters.

### Binning the Integrated Gene Catalogue

We downloaded the non-redundant gene catalogue from Integrated Gene Catalogue (IGC) study^31^ and reads from the associated studies (ERP004605, ERP002061, ERP003612, SRA050230). The reads were mapped using Minimap2 (v.2.2.6) (ref. 48) to the gene catalogue using default parameters and the ‘sr’ prefix for short genomic read mapping. We filtered alignments for a minimum of 95% identity and removed all reads with a read mapping quality less than 30. Hereafter the RPKMs were calculated using VAMB as described above and used for input to VAMB and Canopy clustering. For VAMB we trained the network using a single Tesla Quadro M6000 GPU and clustered the resulting latent representation using 16 CPU cores. For Canopy, we ran canopy on the RPKM-abundance matrix with default parameters for 7 days using 32 CPU cores before we stopped the clustering. The clustering of MSPminer was obtained from the supplementary material of the MSPminer study^36^. Note that the MSPminer authors only used 1,267 of the 1,270 samples of IGC, meaning that the MSPminer bins were made using three samples less compared to Canopy and VAMB. To determine recall and precision, we used CheckM^37^, where we defined recall and precision as recall = completeness, and precision = 1 - (contamination / (contamination + 1)). To compare bins between the different binning programs, a pair of bins were considered overlapping if the number of genes present in both (the intersection) was at least half the number of genes in the larger of the two bins.

### Taxonomic annotation of bins

First the IGC genes were assigned a taxonomy by sequence similarity (MEGA-BLAST) to NCBI RefSeq or the *nt* database (both from October 2018). Genes were assigned strain, species, genus, family, order, class, phylum and domain taxonomy if they had 95, 95, 85, 75, 65, 55, 50 and 45 % identity over 80 % of the gene length, respectively. For each gene multiple taxonomies could be assigned at each level. Bins were assigned taxonomy at strain, species, genus, family, order, class, phylum or domain, if 75, 75, 60, 50, 40, 30, 25 or 25 % of their genes were consistently annotated to a specific taxon and no more than 10 % of the remaining genes were assigned an alternative taxonomy at the given level.

## Acknowledgements

S.R. was supported by the Novo Nordisk Foundation grant NNF14CC0001 and the Jorck Foundation Research Award.

## Author contributions

S.R. conceived the study and guided the analysis, J.N.N. and S.R. performed the analyses, J.N.N. wrote the software, C.K.S., J.A., T.N.P., H.B.N. and O.W. provided guidance and input for the analysis, J.N.N. and S.R. wrote the manuscript with contributions from all co-authors, all authors read and approved the last version of the manuscript.

## Code availability

All code can be found at GitHub at https://github.com/jakobnissen/vamb and is freely available under the permissive MIT license.

## Supplementary table legends

**Table S1**: Hyperparameters in hyperparameter optimization. *sseratio* and *errorsum* corresponds to the VAE parameters α and β, respectively. The “tentative optimum” column gives the apparent optimal value after the first round of optimization, the “final value” shows the chosen value after the second round. See also figures S3-S5.

**Table S2**: Number of recovered bins with more than 200,000 basepairs at or above various precision and recall threshold values when running VAMB with different inputs. For each of the three datasets MetaHIT, CAMI High and CAMI Airways, each of the combinations TNF only, abundance only, or both was tried with autoencoding and without, for a total of 18 different runs. See also Figure S6.

**Table S3**: Number of recovered bins with more than 200,000 basepairs at or above various precision and recall threshold values when running the four binners VAMB, Canopy, METABAT2 and MaxBin on the three datasets MetaHIT, CAMI High and CAMI Airways each. See also Figure 3.

**Table S4:** Binning statistics when binning the Integrated Gene Catalogue using VAMB, MSPminer and Canopy. The 4th to 11th row display the number of recovered bins in large bins (≥ 500 genes) at or above the given recall (R) and precision (P) threshold, as determined by CheckM.

**Table S5:** Various summary statistics of all large (≥ 500 genes) VAMB bins from binning the IGC dataset, and their corresponding “common” bin from MSPminer and/or Canopy, if applicable. Two bins were considered common if they had ≥ 50 % reciprocal overlap of bin content. If a VAMB bin is common to multiple bins from the same binner, the bin with the largest overlap is considered the corresponding bin. The table can be found here: https://doi.org/10.6084/m9.figshare.7415822.

## Additional datasets

VAMB bins (fasta) of the IGC dataset, bins equal to or larger than 500 genes. https://doi.org/10.6084/m9.figshare.7415756

VAMB bins (fasta) of the IGC dataset, all bins. https://doi.org/10.6084/m9.figshare.7415786

