## Supplementary material for "Binning microbial genomes using deep learning"

### Binning microbial genomes using deep learning
Supplementary material

Jakob Nybo Nissen[1],
Casper Kaae Sønderby[2],
Jose Juan Almagro Armenteros[1],
Christopher Heje Grønbech[3,4],
Henrik Bjørn Nielsen[5]
Thomas Nordahl Petersen[6],
Ole Winther[2,3,4],
Simon Rasmussen[7]

December 7, 2018

Contact information:



[1] Department of Bio and Health Informatics, Technical University of Denmark, Kongens Lyngby, Denmark
[2] Bioinformatics Centre, Department of Biology, University of Copenhagen, Copenhagen N, Denmark
[3] Department of Applied Mathematics and Computer Science, Technical University of Denmark, Kongens Lyngby, Denmark
[4] Center for Genomic Medicine, Copenhagen University Hospital, Copenhagen, Denmark
[5] Clinical-Microbiomics A/S, Copenhagen, Denmark
[6] National Food Institute, Technical University of Denmark, Kongens Lyngby, Denmark
[7] Novo Nordisk Foundation Center for Protein Research, Faculty of Health and Medical Sciences, University of Copenhagen, Copenhagen N, Denmark

---

The source code for VAMB can be found at GitHub at https://github.com/jakobnissen/vamb and is freely available under the permissive MIT licence. For the source code creating the plots in this paper, ask us.

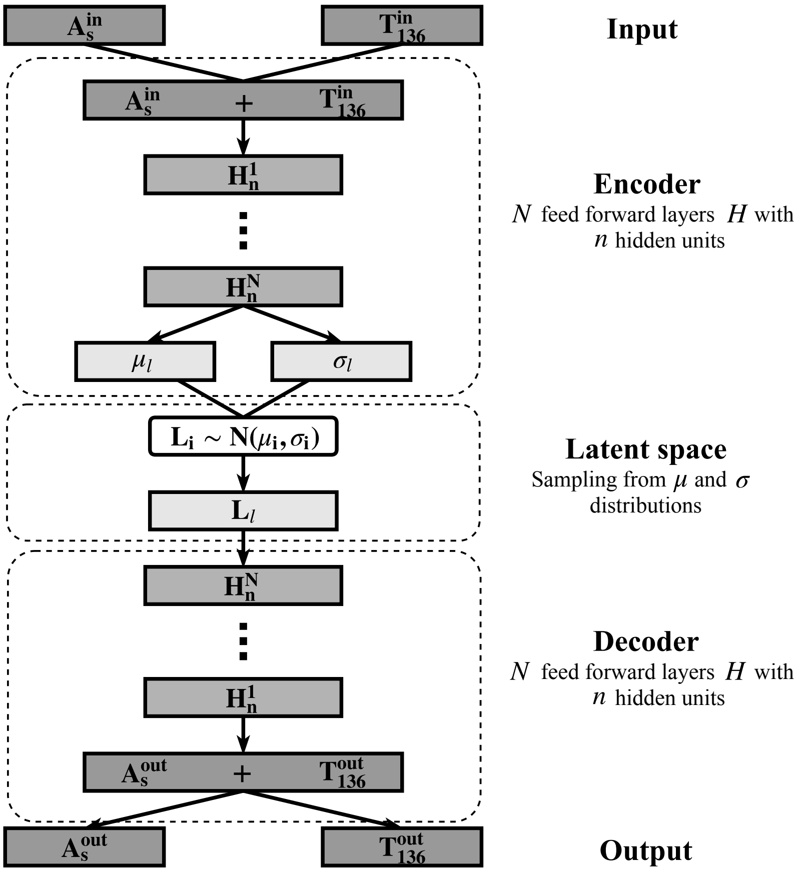


Figure S1: Architecture of VAMB's variational autoencoder. From top to bottom: Input consists of an abundance vector of size $s$ (top left), and a TNF vector of size 136 (top right). These are concatenated to an $s + 136$ vector, then fed through the fully connected encoding layers $H_1$ to $H_N$, each with $n$ neurons. The last layer passes its output to two different fully connected layers termed $\mu$ and $\sigma$, each with $l$ neurons. These steps comprise the encoder. The latent representation is created by sampling a vector of length $l$ where each element $i$ is drawn from the multivariate gaussian distribution $\mathcal{N}(\mu_i, \sigma_i)$. For the decoder, the latent representation is passed through fully connected layers equal to the encoding hidden layers, but ordered in reverse, from $H_N$ to $H_1$. The result is an $s + 136$-vector which is split to an output abundance vector and an output TNF vector. The output abundance vector is softmaxed before output.

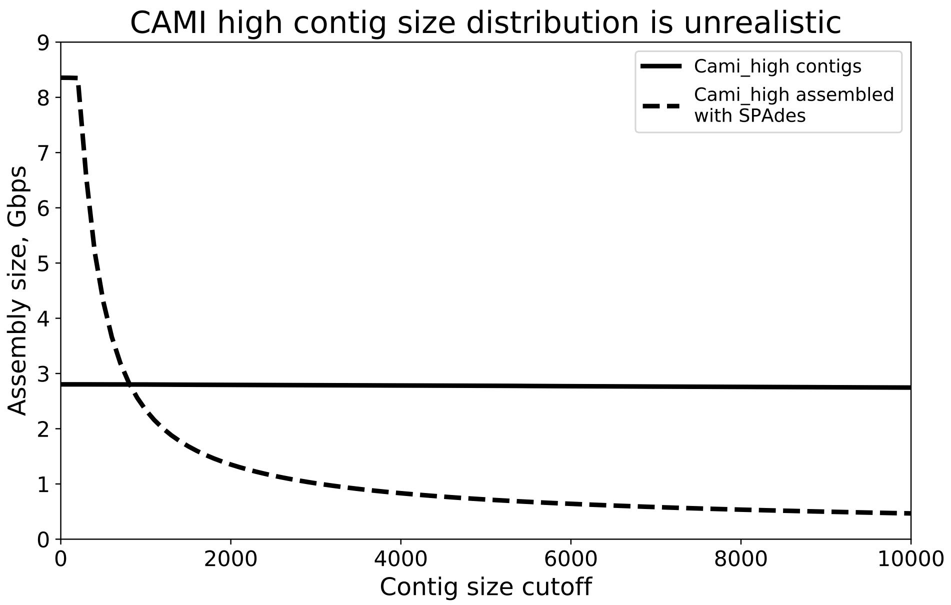


Figure S2: Contig length distribution for CAMI High dataset. On the vertical axis, the total assembly size, i.e. the total number of basepairs in the assembly. On the horizontal axis, the contig size cutoff. Thus, a line crossing the point (4000, $10^9$) means that combined length of all contigs at least 4000 bp in size is $10^9$ bp. In solid, the contig length distribution for the provided contigs of CAMI High, also known as the "gold standard" assembly. The horizontal shape indicates very little of the assembly size is in contigs smaller than 10 kbp. The dotted line is the assembly when assembling the reads with metaSPAdes v. 3.12.0. It is clear by simple inspection that the length distributions are dissimilar, and the metaSPAdes assembly has many more basepairs in smaller contigs.

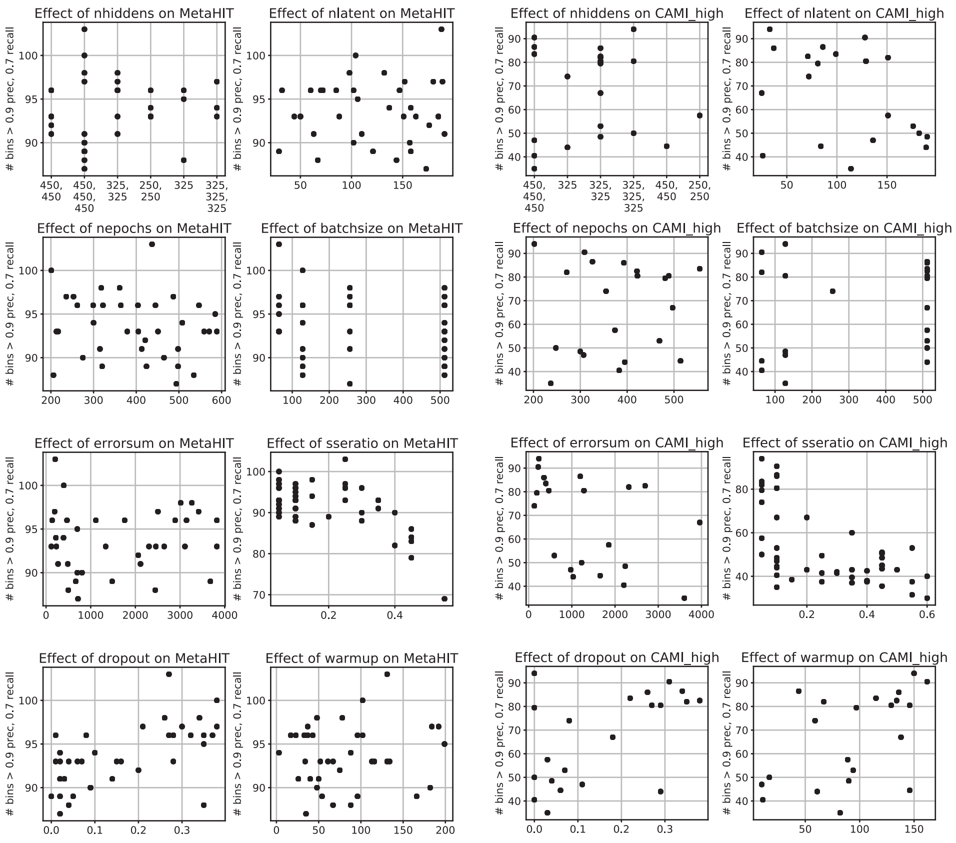


Figure S3: Initial gridsearch for optimal hyperparameters of VAMB. 50 runs for each of MetaHIT and CAMI High was run with hyperparameters randomly chosen within the range displayed on the horizontal axis of the subplots. Note the discrete values for the parameters batchsize (restricted to a power-of-two), and nhiddens, (restricted to the six values shown). The number of decent bins (precision $\geq$ 0.9, recall $\geq$ 0.7) is shown on the vertical axis. Because of the very clear confounding effect of the parameter sseratio, for all subplots except the sseratio subplot, runs with poor values ($\geq$0.3, $\geq$0.2 for MetaHIT and CAMI High) of mseratio has been elided from the plots. The parameters *sseratio* and *capacity* correspond to the parameters $\alpha$ and $\beta$ in the main manuscript, respectively.

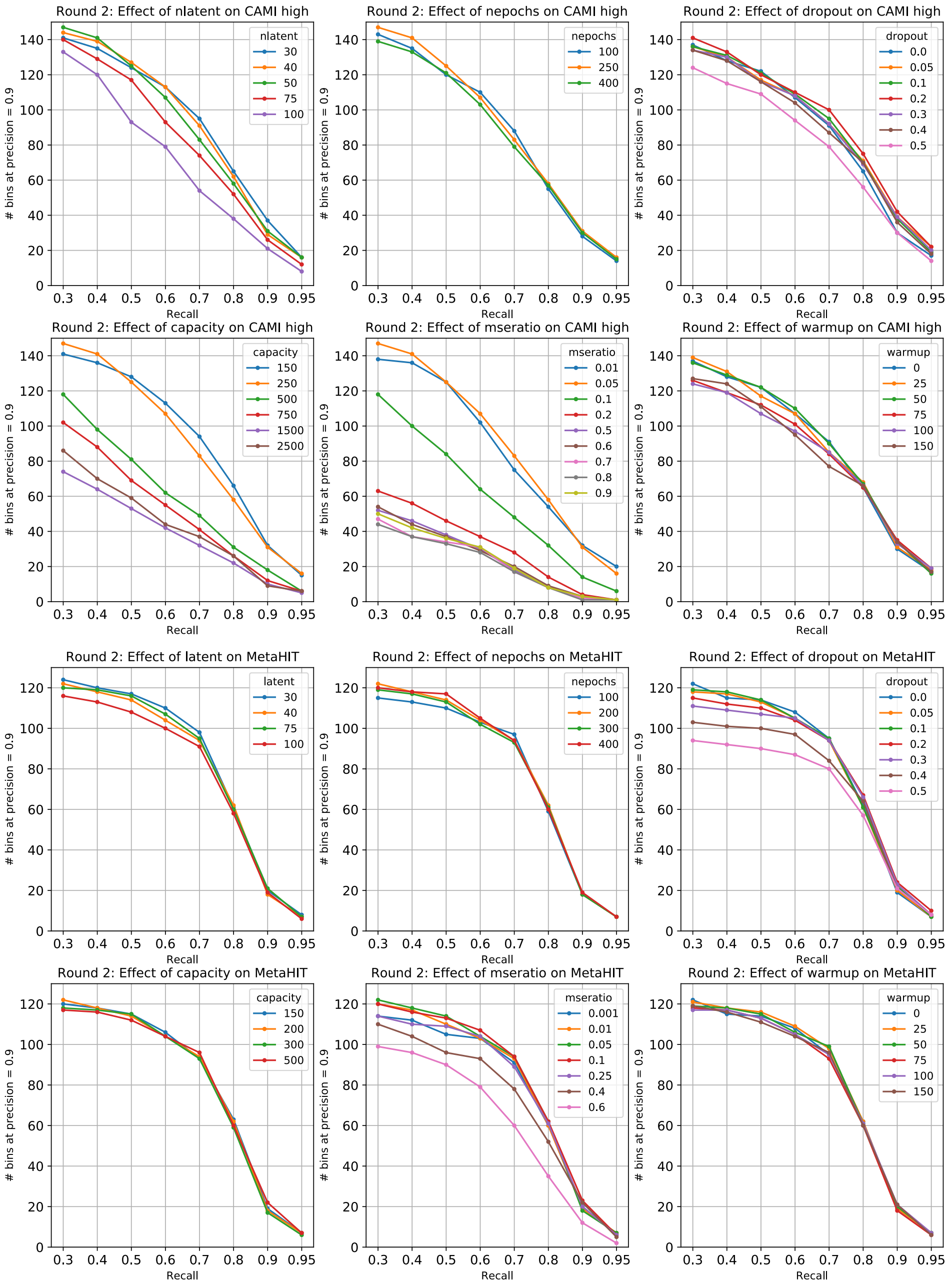


Figure S4: Second round of search for optimal VAMB hyperparameters. In these runs, all hyperparameters except one stay fixed to minimize confounding factors. The default parameters were nhiddens=(325, 325), nlatent=50, nepochs=250, batchsize=128, $\alpha$=0.05, $\beta$=250, dropout=0, warmup=0. On the horizontal axis: Recall threshold value. Vertical axis: Number of recovered bins at precision $\geq 0.9$ at that particular recall threshold value.

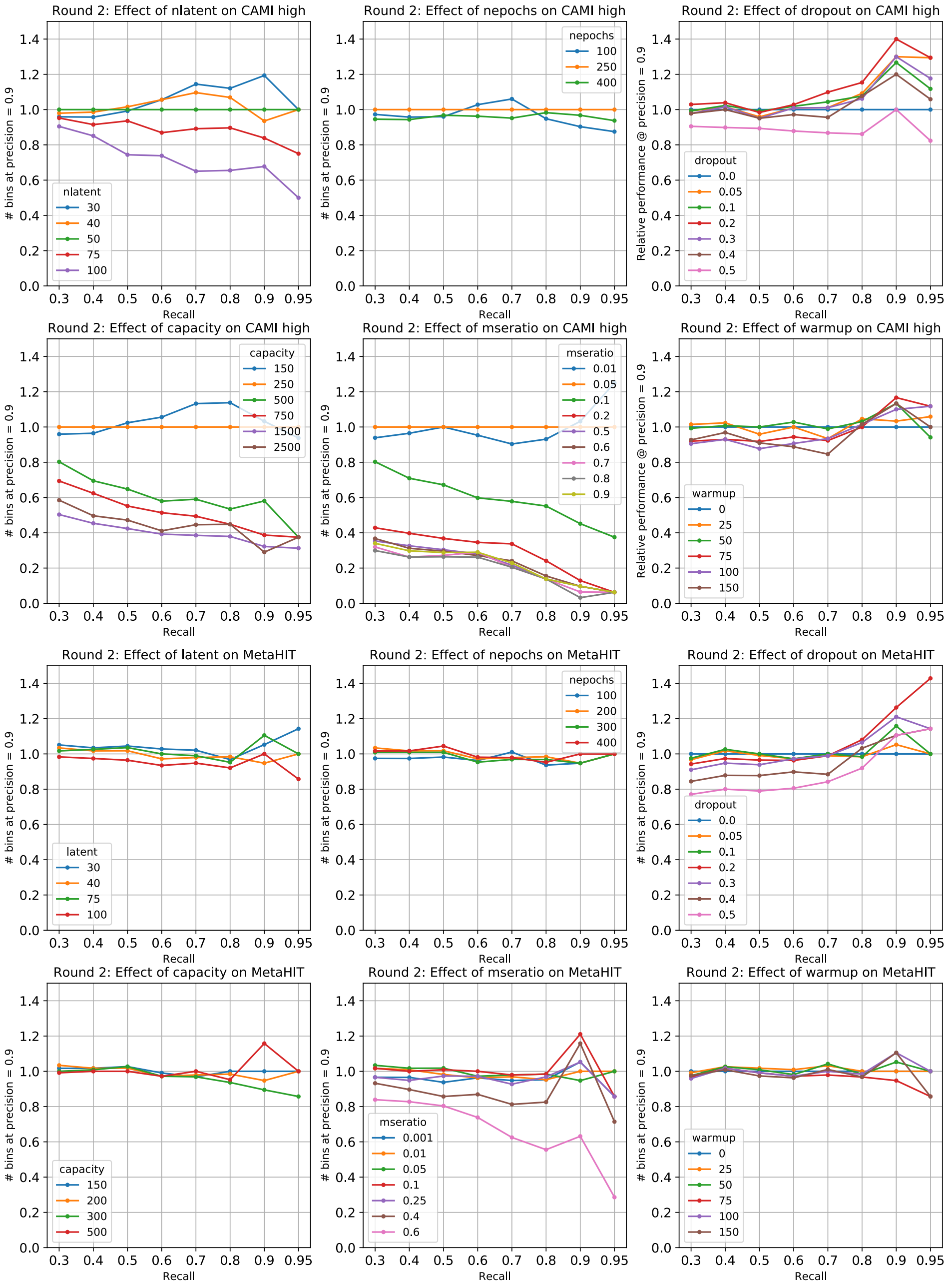


Figure S5: Same as Figure S4, except with values shows as relative to the default parameters.

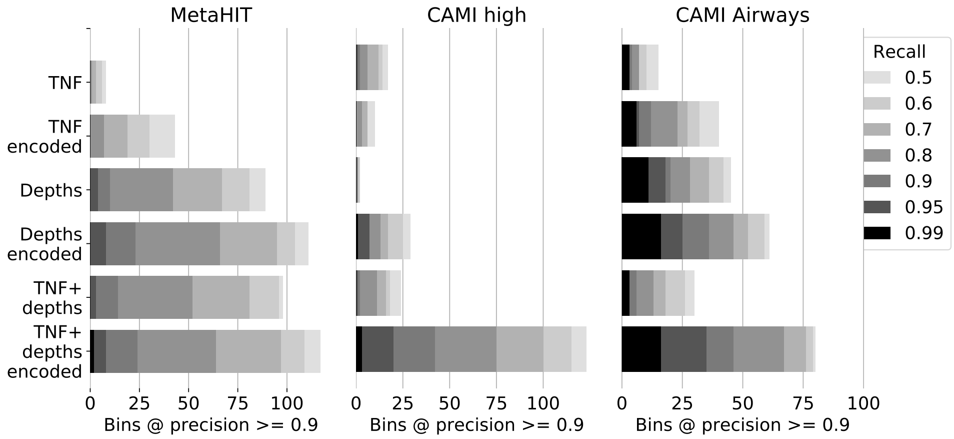


Figure S6: Number of recovered bins for VAMB at precision ≥0.9 and various recall threshold values for the two training datasets MetaHIT and CAMI High, as well at the test dataset CAMI Airways. On the vertical axis, one bar is shown for each input data type: TNF for only TNF data, Depths for only abundance data, and TNF+depths for concatenated TNF and abundance data (VAMB's default), as well as each of those three input types encoded, i.e. where the latent encoding is clustered rather than clustering the raw input data. For all input types except TNF with Cami High, encoding improves the result, leading to more recovered bins.

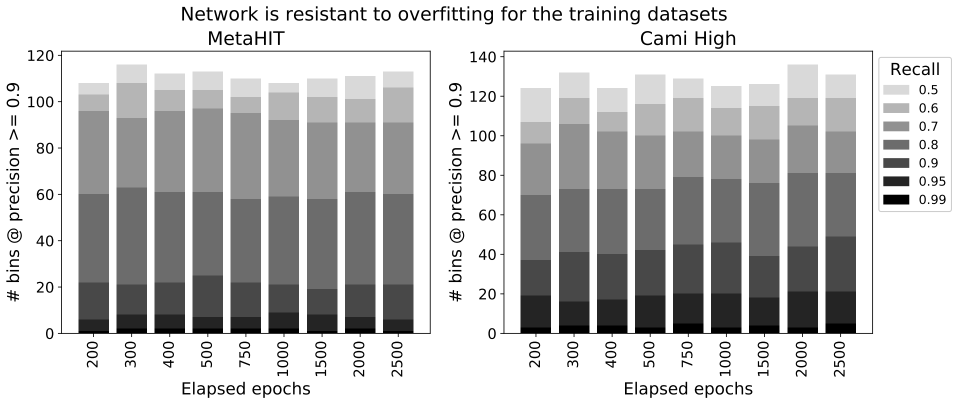


Figure S7: Number of recovered bins at precision ≥0.9 by VAMB at various recall thresholds after training for up to 2,500 epochs. Slight underfitting is seen at 200 epochs, otherwise performance seem to fluctuate randomly, indicating that more epochs does not lead to overfitting on these datasets. Note that overfitting can still occur, for example if the user raises the $\beta$ parameter or increases the number of neurons in the network. The default number of epochs is 500.

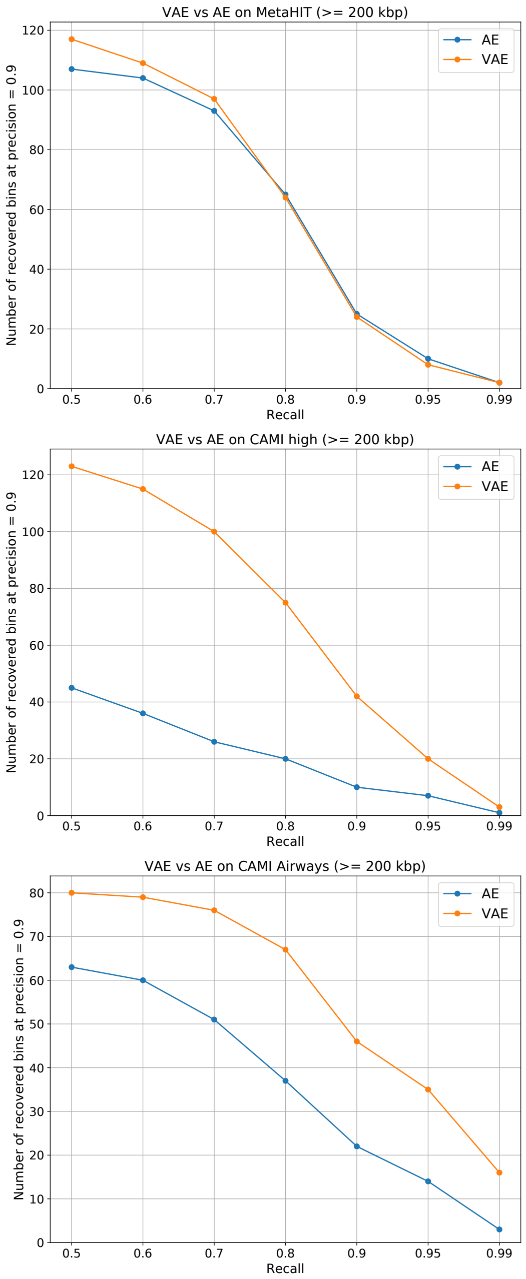


Figure S8: Number of recovered bins on the three benchmark datasets at precision ≥0.9 by VAMB at various recall thresholds when using the default variational autoencoder (VAE, orange line) versus an otherwise identical non-variational autoencoder (AE, blue line)

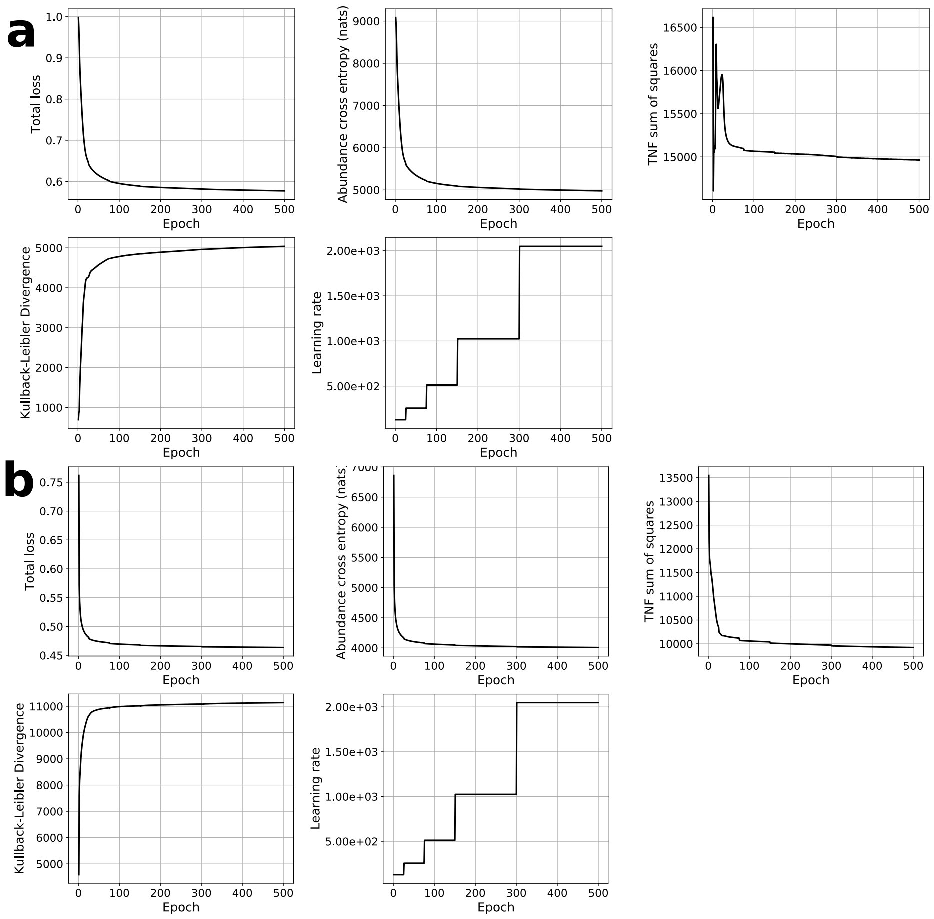


Figure S9: Values of selected training statistics of the VAE when training on the IGC dataset. **a**) Using the default $N_{latent}$ of 40 and dropout of 0.2. Note that TNF sum of squared error does not converge as expected indicating underfitting. **b**: Using $N_{latent}$ of 80 and dropout of 0.0 in order to mitigate underfitting. The latter run is the run used in the main manuscript. In order to be able to learn this very large dataset, the learning rate was decreased from $10^{-3}$ to $10^{-5}$ and starting batch size was decreased from 64 to 128.

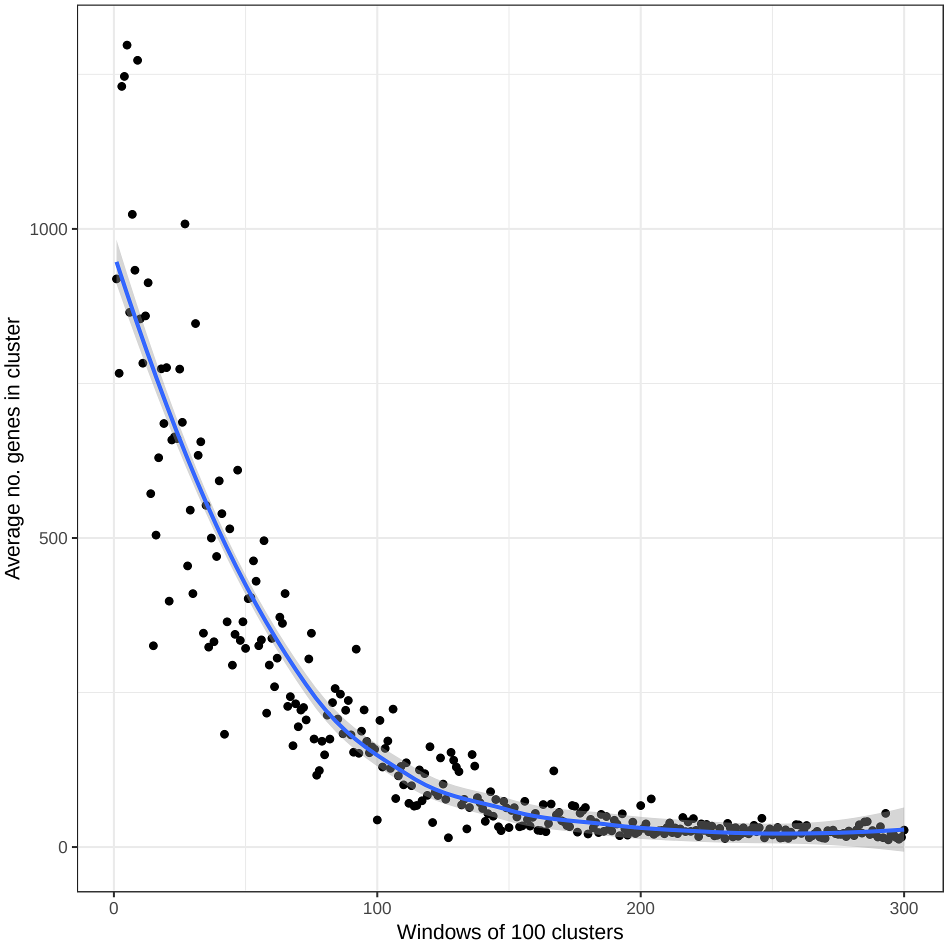


Figure S10: Mean number of genes in the first 30.000 clusters extracted by VAMB from IGC data, calculated across batches of 100 clusters. Since the probability of randomly selecting points from clusters is higher for large clusters than small clusters, the large clusters are sampled first, and few genome-sized clusters are extracted after approximately the first 20.000 clusters.

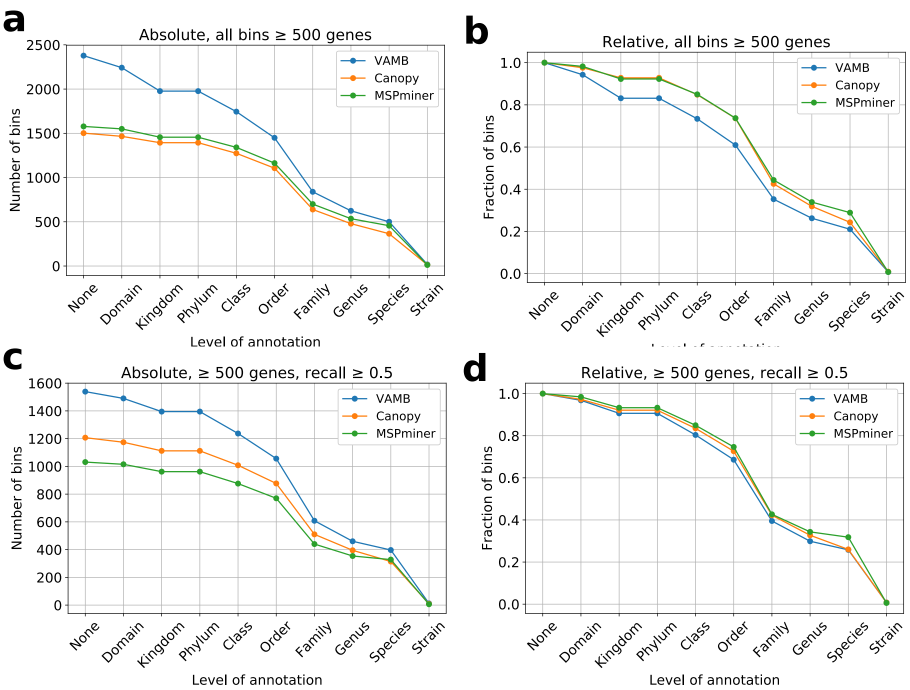


Figure S11: Depth of taxonomic annotation of bins based on database alignment.by sequence similarity (MEGA-BLAST) to NCBI RefSeq or the nt database (both from October 2018). Genes were assigned strain, species, genus, family, order, class, phylum and domain taxonomy if they had 95, 95, 85, 75, 65, 55, 50 and 45 % identity over 80 % of the gene length, respectively. For each gene multiple taxonomies could be assigned at each level. Bins were assigned taxonomy at strain, species, genus, family, order, class, phylum or domain, if 75, 75, 60, 50 40 30, 25 or 20 % of their genes were consistently annotated to a specific taxon and no more than 10 % of the remaining genes were assigned an alternative taxonomy at the given level.

**Algorithm S1** Iterative medoid clustering

```
function CLUSTER(contigs, threshold, N_samples)
    seed ← first contig of contigs
    cluster ← set of all contigs within threshold of seed
    D(cluster) ← mean distance of seed to each other member of cluster
    for trialseed of N_samples randomly sampled contigs from cluster do
        trial ← set of all contigs within threshold of trialseed
        D(trial) ← mean distance of trialseed to each other member of trial
        if D(trial) < D(cluster) then
            seed ← trialseed
            go to 3
        end if
    end for
    output cluster
    contigs ← contigs \ cluster
    if contigs ≠ ∅ then
        go to 2
    end if
end function
```

Supplementary Notes: Iterative medoid clustering algorithm used in the clustering step of VAMB.

Table S1: Hyperparameter search tentative and final optima

| Hyperparameter | Sampled from | Tentative optimum | Final value |
|---|---|---|---|
| nhiddens | ((325,325), (250,250), (450, 450), (325, 325, 325), (325), (450, 450, 450)) | (325, 325) | (325, 325) |
| nlatent | [25, 200] | 50 | 40 |
| nepochs | [200, 600] | 250 | 500 |
| batchsize | (64, 128, 526, 512) | 128 | 64 |
| alpha | [0.05, 0.6] | 0.05 | 0.05 |
| beta | [100, 4000] | 250 | 200 |
| dropout | [0.01, 0.39] | 0 | 0 |
| warmup | [0, 199] | 0 | 0 |

Table S2: Performance of clustering differing input data types

TNF+depths (encoded) on MetaHIT (≥ 200 kbp)

| | Recall | | | | | | |
|---|---|---|---|---|---|---|---|
| Precision | 0.5 | 0.6 | 0.7 | 0.8 | 0.9 | 0.95 | 0.99 |
| 0.7 | 125 | 117 | 101 | 66 | 25 | 8 | 2 |
| 0.8 | 122 | 114 | 100 | 65 | 25 | 8 | 2 |
| 0.9 | 117 | 109 | 97 | 64 | 24 | 8 | 2 |
| 0.95 | 109 | 104 | 92 | 60 | 22 | 7 | 1 |
| 0.99 | 80 | 78 | 69 | 49 | 20 | 5 | 0 |

TNF+depths on MetaHIT (≥ 200 kbp)

| | Recall | | | | | | |
|---|---|---|---|---|---|---|---|
| Precision | 0.5 | 0.6 | 0.7 | 0.8 | 0.9 | 0.95 | 0.99 |
| 0.7 | 112 | 108 | 88 | 56 | 17 | 6 | 2 |
| 0.8 | 104 | 101 | 85 | 54 | 15 | 4 | 0 |
| 0.9 | 98 | 96 | 81 | 52 | 14 | 3 | 0 |
| 0.95 | 89 | 87 | 74 | 49 | 12 | 2 | 0 |
| 0.99 | 65 | 63 | 54 | 38 | 9 | 2 | 0 |

Depths (encoded) on MetaHIT (≥ 200 kbp)

| | Recall | | | | | | |
|---|---|---|---|---|---|---|---|
| Precision | 0.5 | 0.6 | 0.7 | 0.8 | 0.9 | 0.95 | 0.99 |
| 0.7 | 118 | 109 | 98 | 68 | 23 | 8 | 0 |
| 0.8 | 116 | 108 | 97 | 67 | 23 | 8 | 0 |
| 0.9 | 111 | 104 | 95 | 66 | 23 | 8 | 0 |
| 0.95 | 103 | 97 | 89 | 61 | 20 | 7 | 0 |
| 0.99 | 77 | 71 | 64 | 44 | 16 | 3 | 0 |

Depths on MetaHIT (≥ 200 kbp)

| | Recall | | | | | | |
|---|---|---|---|---|---|---|---|
| Precision | 0.5 | 0.6 | 0.7 | 0.8 | 0.9 | 0.95 | 0.99 |
| 0.7 | 97 | 88 | 72 | 44 | 11 | 4 | 0 |
| 0.8 | 94 | 86 | 70 | 43 | 11 | 4 | 0 |
| 0.9 | 89 | 81 | 67 | 42 | 10 | 4 | 0 |
| 0.95 | 81 | 73 | 61 | 36 | 8 | 2 | 0 |
| 0.99 | 61 | 55 | 47 | 29 | 6 | 1 | 0 |

TNF (encoded) on MetaHIT (≥ 200 kbp)

| | Recall | | | | | | |
|---|---|---|---|---|---|---|---|
| Precision | 0.5 | 0.6 | 0.7 | 0.8 | 0.9 | 0.95 | 0.99 |
| 0.7 | 50 | 36 | 23 | 8 | 1 | 0 | 0 |
| 0.8 | 48 | 34 | 22 | 8 | 1 | 0 | 0 |
| 0.9 | 43 | 30 | 19 | 7 | 0 | 0 | 0 |
| 0.95 | 39 | 28 | 18 | 7 | 0 | 0 | 0 |
| 0.99 | 23 | 18 | 12 | 5 | 0 | 0 | 0 |

TNF on MetaHIT (≥ 200 kbp)

| | Recall | | | | | | |
|---|---|---|---|---|---|---|---|
| Precision | 0.5 | 0.6 | 0.7 | 0.8 | 0.9 | 0.95 | 0.99 |
| 0.7 | 14 | 10 | 5 | 3 | 1 | 0 | 0 |
| 0.8 | 11 | 8 | 5 | 3 | 1 | 0 | 0 |
| 0.9 | 8 | 6 | 3 | 1 | 0 | 0 | 0 |
| 0.95 | 7 | 5 | 3 | 1 | 0 | 0 | 0 |
| 0.99 | 4 | 3 | 2 | 1 | 0 | 0 | 0 |

TNF+depths (encoded) on CAMI High (≥ 200 kbp)

| Precision | Recall 0.5 | 0.6 | 0.7 | 0.8 | 0.9 | 0.95 | 0.99 |
|---|---|---|---|---|---|---|---|
| 0.7 | 138 | 126 | 110 | 83 | 47 | 23 | 3 |
| 0.8 | 133 | 123 | 107 | 81 | 46 | 22 | 3 |
| 0.9 | 123 | 115 | 100 | 75 | 42 | 20 | 3 |
| 0.95 | 104 | 98 | 85 | 61 | 35 | 17 | 2 |
| 0.99 | 74 | 71 | 63 | 47 | 28 | 14 | 0 |

TNF+depths on CAMI High (≥ 200 kbp)

| Precision | Recall 0.5 | 0.6 | 0.7 | 0.8 | 0.9 | 0.95 | 0.99 |
|---|---|---|---|---|---|---|---|
| 0.7 | 33 | 25 | 23 | 16 | 5 | 1 | 0 |
| 0.8 | 29 | 22 | 20 | 14 | 3 | 1 | 0 |
| 0.9 | 24 | 18 | 16 | 11 | 2 | 1 | 0 |
| 0.95 | 20 | 15 | 13 | 9 | 2 | 1 | 0 |
| 0.99 | 14 | 12 | 12 | 8 | 2 | 1 | 0 |

Depths (encoded) on CAMI High (≥ 200 kbp)

| Precision | Recall 0.5 | 0.6 | 0.7 | 0.8 | 0.9 | 0.95 | 0.99 |
|---|---|---|---|---|---|---|---|
| 0.7 | 52 | 43 | 32 | 28 | 14 | 11 | 2 |
| 0.8 | 39 | 32 | 23 | 19 | 9 | 9 | 1 |
| 0.9 | 29 | 25 | 17 | 13 | 7 | 7 | 1 |
| 0.95 | 22 | 20 | 14 | 11 | 7 | 7 | 1 |
| 0.99 | 7 | 6 | 3 | 2 | 2 | 2 | 0 |

Depths on CAMI High (≥ 200 kbp)

| Precision | Recall 0.5 | 0.6 | 0.7 | 0.8 | 0.9 | 0.95 | 0.99 |
|---|---|---|---|---|---|---|---|
| 0.7 | 20 | 16 | 13 | 12 | 5 | 2 | 1 |
| 0.8 | 10 | 7 | 6 | 5 | 4 | 2 | 1 |
| 0.9 | 2 | 2 | 1 | 1 | 1 | 0 | 0 |
| 0.95 | 0 | 0 | 0 | 0 | 0 | 0 | 0 |
| 0.99 | 0 | 0 | 0 | 0 | 0 | 0 | 0 |

TNF (encoded) on CAMI High (≥ 200 kbp)

| Precision | Recall 0.5 | 0.6 | 0.7 | 0.8 | 0.9 | 0.95 | 0.99 |
|---|---|---|---|---|---|---|---|
| 0.7 | 20 | 15 | 9 | 4 | 0 | 0 | 0 |
| 0.8 | 18 | 13 | 8 | 4 | 0 | 0 | 0 |
| 0.9 | 10 | 6 | 6 | 3 | 0 | 0 | 0 |
| 0.95 | 6 | 4 | 4 | 2 | 0 | 0 | 0 |
| 0.99 | 0 | 0 | 0 | 0 | 0 | 0 | 0 |

TNF on CAMI High (≥ 200 kbp)

| Precision | Recall 0.5 | 0.6 | 0.7 | 0.8 | 0.9 | 0.95 | 0.99 |
|---|---|---|---|---|---|---|---|
| 0.7 | 27 | 23 | 20 | 11 | 3 | 1 | 0 |
| 0.8 | 25 | 21 | 19 | 11 | 3 | 1 | 0 |
| 0.9 | 17 | 14 | 12 | 6 | 2 | 1 | 0 |
| 0.95 | 13 | 10 | 9 | 4 | 1 | 0 | 0 |
| 0.99 | 11 | 8 | 7 | 3 | 1 | 0 | 0 |

TNF+depths (encoded) on CAMI Airways (≥ 200 kbp)

Recall

| Precision | 0.5 | 0.6 | 0.7 | 0.8 | 0.9 | 0.95 | 0.99 |
|---|---|---|---|---|---|---|---|
| 0.7 | 97 | 95 | 91 | 80 | 53 | 42 | 18 |
| 0.8 | 90 | 88 | 85 | 76 | 52 | 41 | 18 |
| 0.9 | 80 | 79 | 76 | 67 | 46 | 35 | 16 |
| 0.95 | 61 | 61 | 58 | 52 | 39 | 31 | 14 |
| 0.99 | 37 | 37 | 34 | 31 | 27 | 20 | 11 |

TNF+depths on CAMI Airways (≥ 200 kbp)

Recall

| Precision | 0.5 | 0.6 | 0.7 | 0.8 | 0.9 | 0.95 | 0.99 |
|---|---|---|---|---|---|---|---|
| 0.7 | 39 | 33 | 24 | 17 | 8 | 4 | 3 |
| 0.8 | 35 | 31 | 22 | 15 | 8 | 4 | 3 |
| 0.9 | 30 | 26 | 18 | 13 | 6 | 3 | 3 |
| 0.95 | 27 | 23 | 16 | 11 | 5 | 3 | 3 |
| 0.99 | 17 | 13 | 10 | 8 | 4 | 2 | 2 |

Depths (encoded) on CAMI Airways (≥ 200 kbp)

Recall

| Precision | 0.5 | 0.6 | 0.7 | 0.8 | 0.9 | 0.95 | 0.99 |
|---|---|---|---|---|---|---|---|
| 0.7 | 75 | 70 | 60 | 52 | 39 | 28 | 18 |
| 0.8 | 70 | 67 | 57 | 51 | 39 | 28 | 18 |
| 0.9 | 61 | 59 | 52 | 46 | 36 | 25 | 16 |
| 0.95 | 52 | 50 | 45 | 39 | 31 | 21 | 14 |
| 0.99 | 37 | 36 | 32 | 28 | 23 | 18 | 12 |

Depths on CAMI Airways (≥ 200 kbp)

Recall

| Precision | 0.5 | 0.6 | 0.7 | 0.8 | 0.9 | 0.95 | 0.99 |
|---|---|---|---|---|---|---|---|
| 0.7 | 53 | 49 | 43 | 33 | 23 | 20 | 11 |
| 0.8 | 49 | 45 | 39 | 30 | 22 | 19 | 11 |
| 0.9 | 45 | 42 | 36 | 28 | 20 | 18 | 11 |
| 0.95 | 40 | 37 | 33 | 26 | 19 | 17 | 10 |
| 0.99 | 26 | 25 | 23 | 18 | 13 | 12 | 6 |

TNF (encoded) on CAMI Airways (≥ 200 kbp)

Recall

| Precision | 0.5 | 0.6 | 0.7 | 0.8 | 0.9 | 0.95 | 0.99 |
|---|---|---|---|---|---|---|---|
| 0.7 | 47 | 38 | 33 | 29 | 16 | 10 | 7 |
| 0.8 | 41 | 33 | 28 | 24 | 13 | 7 | 6 |
| 0.9 | 40 | 32 | 27 | 23 | 12 | 7 | 6 |
| 0.95 | 32 | 28 | 23 | 20 | 10 | 6 | 6 |
| 0.99 | 26 | 23 | 19 | 18 | 9 | 5 | 5 |

Table S3: Performance of binner on benchmark datasets

Canopy on CAMI Airways (bins ≥ 200 kbp)

| | Recall | | | | | | |
|---|---|---|---|---|---|---|---|
| Precision | 0.5 | 0.6 | 0.7 | 0.8 | 0.9 | 0.95 | 0.99 |
| 0.7 | 12 | 8 | 4 | 3 | 2 | 2 | 0 |
| 0.8 | 10 | 7 | 4 | 3 | 2 | 2 | 0 |
| 0.9 | 5 | 4 | 3 | 2 | 1 | 1 | 0 |
| 0.95 | 1 | 0 | 0 | 0 | 0 | 0 | 0 |
| 0.99 | 0 | 0 | 0 | 0 | 0 | 0 | 0 |

MaxBin on CAMI Airways (bins ≥ 200 kbp)

| | Recall | | | | | | |
|---|---|---|---|---|---|---|---|
| Precision | 0.5 | 0.6 | 0.7 | 0.8 | 0.9 | 0.95 | 0.99 |
| 0.7 | 58 | 53 | 43 | 32 | 14 | 8 | 3 |
| 0.8 | 51 | 46 | 37 | 29 | 12 | 7 | 2 |
| 0.9 | 39 | 37 | 29 | 25 | 10 | 6 | 2 |
| 0.95 | 32 | 31 | 25 | 21 | 9 | 6 | 2 |
| 0.99 | 14 | 13 | 12 | 11 | 6 | 5 | 2 |

METABAT2 on CAMI Airways (bins ≥ 200 kbp)

| | Recall | | | | | | |
|---|---|---|---|---|---|---|---|
| Precision | 0.5 | 0.6 | 0.7 | 0.8 | 0.9 | 0.95 | 0.99 |
| 0.7 | 109 | 94 | 68 | 50 | 27 | 18 | 8 |
| 0.8 | 105 | 90 | 66 | 49 | 27 | 18 | 8 |
| 0.9 | 97 | 84 | 63 | 48 | 26 | 18 | 8 |
| 0.95 | 88 | 76 | 57 | 45 | 24 | 17 | 7 |
| 0.99 | 65 | 60 | 44 | 35 | 20 | 15 | 7 |

VAMB on CAMI Airways (bins ≥ 200 kbp)

| | Recall | | | | | | |
|---|---|---|---|---|---|---|---|
| Precision | 0.5 | 0.6 | 0.7 | 0.8 | 0.9 | 0.95 | 0.99 |
| 0.7 | 97 | 95 | 91 | 80 | 53 | 42 | 18 |
| 0.8 | 90 | 88 | 85 | 76 | 52 | 41 | 18 |
| 0.9 | 80 | 79 | 76 | 67 | 46 | 35 | 16 |
| 0.95 | 61 | 61 | 58 | 52 | 39 | 31 | 14 |
| 0.99 | 37 | 37 | 34 | 31 | 27 | 20 | 11 |

Canopy on CAMI high (bins ≥ 200 kbp)

| | Recall | | | | | | |
|---|---|---|---|---|---|---|---|
| Precision | 0.5 | 0.6 | 0.7 | 0.8 | 0.9 | 0.95 | 0.99 |
| 0.7 | 0 | 0 | 0 | 0 | 0 | 0 | 0 |
| 0.8 | 0 | 0 | 0 | 0 | 0 | 0 | 0 |
| 0.9 | 0 | 0 | 0 | 0 | 0 | 0 | 0 |
| 0.95 | 0 | 0 | 0 | 0 | 0 | 0 | 0 |
| 0.99 | 0 | 0 | 0 | 0 | 0 | 0 | 0 |

MaxBin on CAMI high (bins ≥ 200 kbp)

| | Recall | | | | | | |
|---|---|---|---|---|---|---|---|
| Precision | 0.5 | 0.6 | 0.7 | 0.8 | 0.9 | 0.95 | 0.99 |
| 0.7 | 24 | 19 | 15 | 9 | 1 | 1 | 1 |
| 0.8 | 22 | 17 | 13 | 8 | 1 | 1 | 1 |
| 0.9 | 16 | 12 | 9 | 5 | 1 | 1 | 1 |
| 0.95 | 9 | 6 | 5 | 2 | 1 | 1 | 1 |
| 0.99 | 5 | 2 | 2 | 1 | 0 | 0 | 0 |

METABAT2 on CAMI high (bins ≥ 200 kbp)

| Precision | Recall 0.5 | 0.6 | 0.7 | 0.8 | 0.9 | 0.95 | 0.99 |
|---|---|---|---|---|---|---|---|
| 0.7 | 97 | 87 | 71 | 45 | 20 | 10 | 2 |
| 0.8 | 94 | 86 | 70 | 44 | 19 | 10 | 2 |
| 0.9 | 87 | 79 | 64 | 40 | 16 | 7 | 1 |
| 0.95 | 74 | 68 | 55 | 32 | 11 | 4 | 1 |
| 0.99 | 34 | 32 | 28 | 19 | 8 | 4 | 1 |

VAMB on CAMI high (bins ≥ 200 kbp)

| Precision | Recall 0.5 | 0.6 | 0.7 | 0.8 | 0.9 | 0.95 | 0.99 |
|---|---|---|---|---|---|---|---|
| 0.7 | 138 | 126 | 110 | 83 | 47 | 23 | 3 |
| 0.8 | 133 | 123 | 107 | 81 | 46 | 22 | 3 |
| 0.9 | 123 | 115 | 100 | 75 | 42 | 20 | 3 |
| 0.95 | 104 | 98 | 85 | 61 | 35 | 17 | 2 |
| 0.99 | 74 | 71 | 63 | 47 | 28 | 14 | 0 |

Canopy on MetaHIT (bins ≥ 200 kbp)

| Precision | Recall 0.5 | 0.6 | 0.7 | 0.8 | 0.9 | 0.95 | 0.99 |
|---|---|---|---|---|---|---|---|
| 0.7 | 96 | 89 | 66 | 26 | 4 | 1 | 0 |
| 0.8 | 93 | 86 | 64 | 25 | 4 | 1 | 0 |
| 0.9 | 84 | 79 | 60 | 24 | 3 | 0 | 0 |
| 0.95 | 77 | 73 | 55 | 22 | 3 | 0 | 0 |
| 0.99 | 66 | 63 | 48 | 20 | 3 | 0 | 0 |

MaxBin on MetaHIT (bins ≥ 200 kbp)

| Precision | Recall 0.5 | 0.6 | 0.7 | 0.8 | 0.9 | 0.95 | 0.99 |
|---|---|---|---|---|---|---|---|
| 0.7 | 50 | 41 | 23 | 10 | 4 | 0 | 0 |
| 0.8 | 35 | 29 | 17 | 8 | 2 | 0 | 0 |
| 0.9 | 23 | 21 | 14 | 6 | 1 | 0 | 0 |
| 0.95 | 14 | 12 | 10 | 5 | 1 | 0 | 0 |
| 0.99 | 4 | 4 | 4 | 2 | 0 | 0 | 0 |

METABAT2 on MetaHIT (bins ≥ 200 kbp)

| Precision | Recall 0.5 | 0.6 | 0.7 | 0.8 | 0.9 | 0.95 | 0.99 |
|---|---|---|---|---|---|---|---|
| 0.7 | 102 | 91 | 65 | 31 | 4 | 3 | 0 |
| 0.8 | 99 | 88 | 62 | 31 | 4 | 3 | 0 |
| 0.9 | 89 | 82 | 58 | 29 | 3 | 2 | 0 |
| 0.95 | 83 | 78 | 56 | 28 | 3 | 2 | 0 |
| 0.99 | 58 | 55 | 41 | 22 | 3 | 2 | 0 |

VAMB on MetaHIT (bins ≥ 200 kbp)

| Precision | Recall 0.5 | 0.6 | 0.7 | 0.8 | 0.9 | 0.95 | 0.99 |
|---|---|---|---|---|---|---|---|
| 0.7 | 125 | 117 | 101 | 66 | 25 | 8 | 2 |
| 0.8 | 122 | 114 | 100 | 65 | 25 | 8 | 2 |
| 0.9 | 117 | 109 | 97 | 64 | 24 | 8 | 2 |
| 0.95 | 109 | 104 | 92 | 60 | 22 | 7 | 1 |
| 0.99 | 80 | 78 | 69 | 49 | 20 | 5 | 0 |

Table S4: Summary statistics for binning on IGC dataset

| Binner | Canopy | MSPminer | VAMB |
|---|---|---|---|
| Genes clustered (m) | 3.86 | 3.36 | 5.46 |
| Large bins | 1578 | 1502 | 2378 |
| Genes in large bins (m) | 2.71 | 3.31 | 4.60 |
| Prec. $\geq$ 0.9, Rec. $\geq$ 0.5 | 1002 | 1095 | 1277 |
| Prec. $\geq$ 0.9, Rec. $\geq$ 0.6 | 923 | 996 | 1169 |
| Prec. $\geq$ 0.9, Rec. $\geq$ 0.7 | 822 | 904 | 1066 |
| Prec. $\geq$ 0.9, Rec. $\geq$ 0.8 | 653 | 760 | 926 |
| Prec. $\geq$ 0.9, Rec. $\geq$ 0.9 | 429 | 558 | 675 |
| Prec. $\geq$ 0.9, Rec. $\geq$ 0.95 | 246 | 344 | 406 |
| Prec. $\geq$ 0.9, Rec. $\geq$ 0.99 | 30 | 55 | 76 |
